# Chaperone quality control in liquid-phase separated organelles

**DOI:** 10.1101/2024.01.23.576883

**Authors:** Tom Scheidt, Edward A. Lemke

## Abstract

Molecular chaperones, central to the cellular proteostasis network, play an essential role in preventing the formation and proliferation of harmful aggregates associated with neurodegenerative diseases. Notably, for many intrinsically disordered proteins (IDPs), which are prone to form such damaging deposits, the formation of nano-clusters and phase separation into organelles prior to aggregation have been observed. The impact of molecular chaperones on such assemblies, remains unclear. In our study, we concentrated on the family of small heat shock proteins (sHsps), which are typically dynamic and form large oligomeric structures. While sHsps are mainly structured/folded proteins, they can undergo transient multivalent interactions, like many IDPs. Thus, sHsps might be a suitable regulator for vital and ubiquitous formation of membrane-less organelles in eukaryotic cells rich in IDPs and to inhibit aberrant aggregation. Here we show, using microfluidic diffusional sizing, that the formation of nano-clusters of FUS, associated with neurodegenerative diseases can be inhibited by the presence of sHsps. Furthermore, we identify that, depending on their assembly state, sHsps are capable of targeting specifically the interface between the dense droplet phase and the dilute phase not only of FUS but also of TDP-43, likely because the interface is the primary starting point for fibril formation or protein aggregation in general. Our findings emphasise the impact of molecular chaperones on maintaining the homeostasis of IDPs in the dilute and condensed phase. This could help to understand how chaperone dysregulation can influence aberrant protein association.

## Introduction

Neurodegenerative diseases, such as Alzheimer’s, Parkinson’s, amyotrophic lateral sclerosis (ALS) and frontotemporal dementia (FTD), have long been associated with the formation of amyloid fibrils^1,2^. This includes a set of proteins, rich in intrinsically disordered regions, such as α-synuclein, Tau, Huntingtin, Fused in Sarcoma (FUS) or TAR DNA-binding protein 43 (TDP-43)^3–6^. Besides the possibility of fibril formation through primary and secondary nucleation in the dilute phase^7–11^, recent studies promote the concept of pathological phase transition, where fibril formation has been observed within the dense liquid phase^12–15^. In fact, the majority of neurodegenerative disease associated proteins have been shown to be recruited to stress granules, ribonucleoprotein (RNP) granules, Cajal bodies, or other phase separated membrane-less compartments^15–17^. Nevertheless, despite their known individual properties to phase separate into highly concentrated droplets *in vitro*, it is uncertain on how their cellular homeostasis is maintained and their aberration/ageing into fibrillar aggregates is prevented.

Physiological mechanisms inhibiting such early pathogenic events can be upregulation of the chaperone or proteostasis machinery and thus reverse excessive homotypic interactions. Similar to the formation of dynamic membrane-less cellular compartments, which are based on multivalent interactions, the class of small heat shock proteins (sHsps) are highly dynamic and multivalent assemblies themselves and the first line of defence against protein unfolding stress^18,19^. The ubiquitous sHsps constitute a diverse chaperone family sharing all a conserved α-crystallin domain (ACD) as the core structural element surrounded by variable disordered N- and C-terminal extensions (NTE/CTE)^20^. Even though the monomeric mass is ranging between 17 and 28 kDa for all 10 members in humans, called HspB1-10, accordingly^21^, these chaperones can form highly polydisperse homo- and heterooligomeric structures. Dimers formed via the β-sandwich structured ACD domain generate the smallest building block, with NTE and CTE allowing for a hierarchical assembly to hollow sphere-like structures of 2-32 units^22^. Even though, half of the human HspBs are ubiquitously expressed, some show higher expression levels in certain cell types, such as neurons, skeletal and cardiac muscle cells. Thus, it is maybe not surprising that mutations in HspBs cause neuropathies, myopathies and cardiopathies^21^. In particular, HspB1, HspB2, HspB3, HspB5, HspB6 and HspB8 are specifically associated with neurodegenerative diseases and for even more, interactions to amyloidogenic proteins, such as TDP-43, Tau, α-synuclein, polyQ, Aβ and SOD-1, and effects on aggregation have been shown^23–27^. Some of these amyloidogenic proteins have been found to enrich in cellular organelles like stress granules but also as solid cellular inclusions^28–30^. The sHsps function primarily as ATP-independent holdases and hence, sequester unfolded proteins and prevent their aggregation and allow refolding^31^. Additionally, sHsps support the cellular degradation system by substrate routing^32,33^, regulating cytoskeleton assembly^34,35^ and protein folding and integrity and are key players in cellular stress response and apoptosis^36,37^. On the other hand, the diverse structural arrangements due to polydisperse assembly, allow sHsps to target a wider range of different molecules to promote their stability^25^.

The view on protein maturation/ageing is controversial. Commonly accepted is the progress of proteins to urge to the most stable thermodynamic state with the lowest free-energy over time, which leads ultimately to amyloid fibril formation. Between such a final conformation and a *de novo* unfolded generated protein monomer are a multitude of local transient or semi-stable free-energy minima. Such transient compositions ranging in size from (partially) folded monomeric proteins to oligomeric assemblies over amorphous nano- or even micellular clusters, tens to hundreds of nanometre, to macroscopic droplets^38,39^. On the one hand, recent investigations show that such macroscopic droplets stabilise individual proteins within the dense state^40,41^, but on the other hand, other experiments show enriched protein aggregation for some systems, especially at the phase boundaries of dense and dilute phase of droplets and organelles, the interface^12,13,15,42^. In cells, with its out-of-equilibrium thermodynamics such transient compositions are highly dynamic and tightly regulated, in particular by molecular chaperones. Our aim is to understand where and how sHsps come into play controlling these transient protein compositions for disordered proteins.

## Results

### Effect of small heat shock proteins on disorder proteins above saturation concentrations

Cellular stress goes hand in hand with the formation of stress granules and increased expression of molecular chaperones. Nevertheless, how the accumulation of proteins into dense droplets and the generation of sHsps are connected or whether both mechanisms work fully independent for cell protection is unclear. In order to study the influence of individual sHsp constitution on IDPs, in particular FUS and TDP-43, above saturation concentration, we mixed various sHsps with FUS or TDP-43, respectively (see Fig. 1 and Supplementary Fig. S1). FUS and TDP-43 were expressed as a fusion construct with MBP, which inhibits protein condensation (see Fig. 1 A and Supplementary Fig. S1 A). The condensation of both IDPs could be initiated by the addition of TEV protease, which induces cleavage of the MBP-tag (see Fig. 1 B and Supplementary Fig. S1). The microscope images and intensity linescans of individual droplets, reveal a preferential accumulation for HspB1, HspB4 and HspB5, at the interface of the IDP droplets, whereas for the core-domain of HspB1, HspB1-CD and HspB6 there is no specific interface accumulation observable. For the latter, we see rather a slight exclusion from the dense phase of the droplets. The exact same effect can also be observed for TDP-43 droplets (see Supplementary Fig. S1). These chaperone specific observations correlate well with the hydrodynamic radii measured using microfluidic diffusional sizing (see Fig. 1 D and Fig. 3 A). The three sHsps HspB1, HspB4 and HspB5 give a radius of 7.9±1.2 nm, 8.1±0.8 nm and 7.6±1.2 nm, respectively. These results align well with values found in the literature and the known formation of large oligomeric constitutions of around 16-28 monomers^24,43–45^. The fusion of monomeric enhanced Green Fluorescent Protein (meGFP) to the sHsps did not appear to alter their normal oligomeric assembly, indicating that meGFP can be utilized as a reliable fusion tag in our studies without influencing the inherent structural characteristics of these proteins. In contrast, HspB1-CD and HspB6 have an average hydrodynamic radius of 2.6±0.4 nm and 3.6±0.2 nm, indicating a monomeric and dimeric constitution, respectively in line with expectations from literature^46^. HspB1-CD was designed to lack the NTD and CTD and thus is not able to oligomerize.

**Fig. 1.**
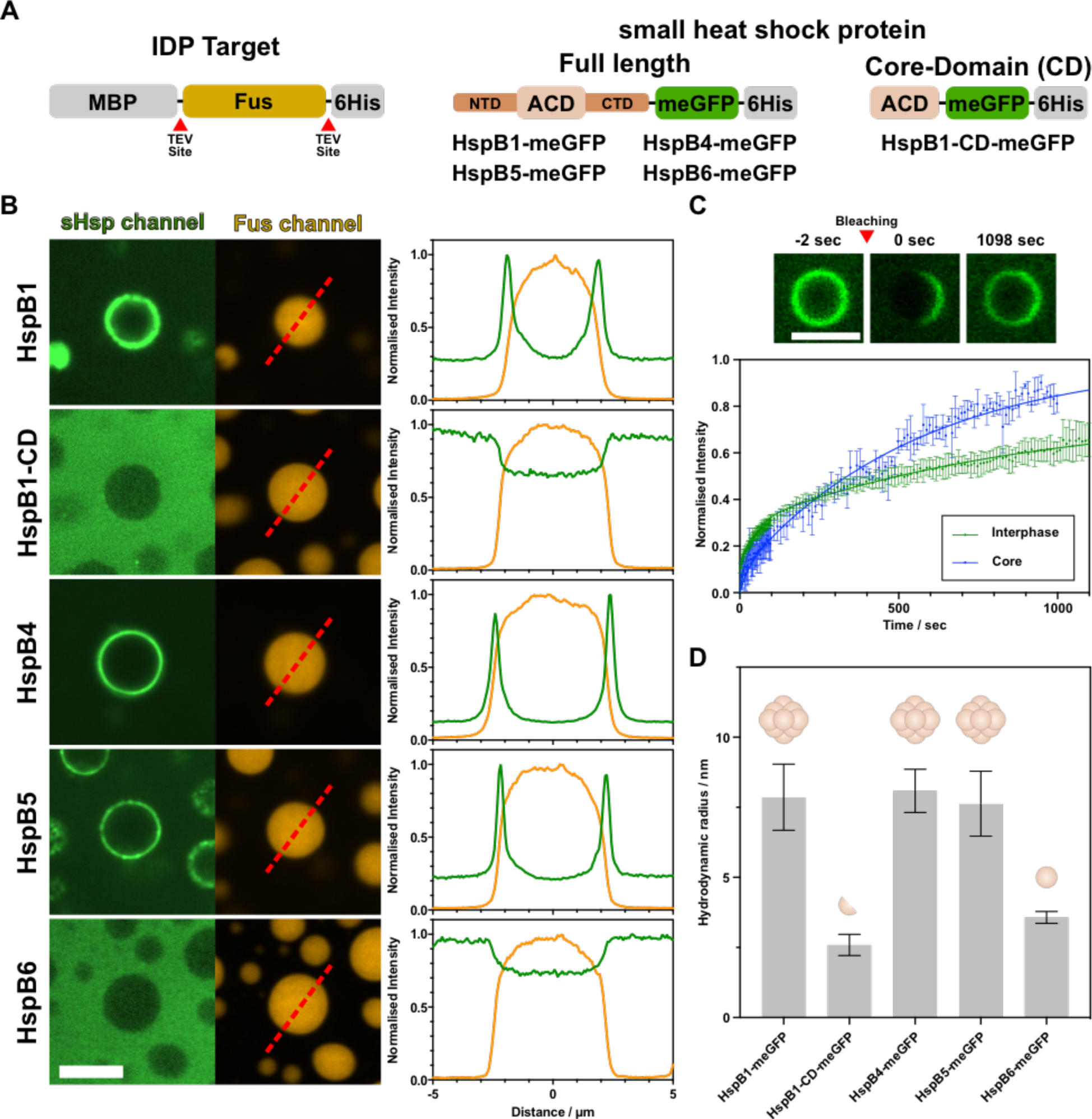
Effect of small heat shock proteins on FUS above saturation conditions. **(A)** Schematic of used protein constructs with FUS as a representative IDP target. Wild-type FUS construct contains an N-terminal maltose-binding protein (MBP) for increased solubility and a C-terminal 6-histidine tag. Both can be cleaved off by TEV protease (left). The sHsps with their N- and C-terminal domains (NTD/CTD) and the α-crystallin domain (ACD) were fused to an meGFP (center). We used also a construct containing only the ACD domain and meGFP (right). **(B)** Spinning disk images of bulk droplet assays with 10 µM FUS and 3 µM HspB1, 5 µM HspB1-CD, 5 µM HspB4, 6 µM HspB5 and 9 µM HspB6 in 10 mM HEPES pH 7.2, 200 mM KCl, 3% glycerol, 1 mM DTT buffer, respectively (left). FUS sample was supplemented with 1:100 MBP-FUS-mscarlet3 and phase separation was triggered by the addition of TEV. Scale bar is 5 µm. On the right, normalised line profiles of the microscope images add the indicated areas (red dashed lines). Scale bar is 5 µm. **(C)** Confocal scans of FRAP of HspB1 in samples similar to **(B)**. Scale bar is 2.5 µm. Normalised fluorescence recovery of multiple droplet rims and centers over time plotted below and fitted to a two-phase association model. **(D)** Hydrodynamic radius measured by diffusional sizing of individual sHsps alone in 20 mM PBS buffer, 0.1% Tween 20, pH 7.4 at 21°C.

For HspB1, which accumulates at the interface, we were measuring the fluorescence recovery after photobleaching (FRAP) of multiple droplets (see Fig. 1 C) and fitted to a two-phase association model. We can observe a slow recovery of chaperone signal at the interface and at the centre of the droplet. However, the initial recovery of HspB1 at the interface (green curve) is faster than inside the droplet (blue curve). The slower recovery of signal inside the droplets compared to the interface is likely caused by the interfacial barrier between dense and dilute phase, restricting the exchange of molecules between the phases. For the interface, the rate constants k_fast_ and k_slow_ are 0.029 sec^-1^ and 0.001 sec^-1^, respectively, with 37.1% fast fraction. The droplet center on the other hand gives rates of k_fast_ and k_slow_ are 0.037 sec^-1^ and 0.002 sec^-1^, respectively, with 8.9% fast fraction. The curves indicating a somewhat dynamic system, allowing the chaperone to diffuse and specifically accumulate at the interface again. In this context, for the interface of condensates it has been recently predicted and found to promote amyloid formation instead of occurring homogeneously inside of droplets^12,13,42^.

In order to observe and characterise protein dynamics within and at the interface of droplet, we made use of fluorescence anisotropy measurements of our molecule of interest. The smaller and/or more mobile the molecule the faster it depolarises polarized excitation light, due to its fast rotation which results in lower anisotropy. In particular, the enrichment of larger, slower rotating FUS molecules, potentially pre-fibrillar, at the droplet interface could be identified using anisotropy measurements (see Fig. 2 B top). Therefore, we excited FUS supplemented (1:500) with FUS-fused mscarlet3 with polarised light and detected the polarization status of the emitted light (see Fig. 2 A). We found high anisotropy indicating that larger FUS complexes or FUS bound to other molecules have a slower rotation compared to their monomeric counterparts. The same idea can be followed exciting meGFP of the sHsps specifically. By thresholding the overall images, we could generate a region of interest for the dense and the dilute phase and access the pixel-by-pixel anisotropy distribution for those regions (see Fig. 2 C) and identify the peak anisotropy (see Table 1), respectively. For the cases where the anisotropy of FUS was measured, it can be observed that the overall anisotropy in the dilute phase is increasing in the presence of sHsps, especially for the HspB1 and HspB4, both known to form larger constitutions (see Fig. 2 C and Tab. 1). Besides the increased anisotropy of the FUS dilute phase, also an increased peak anisotropy in the FUS dense phase can be observed for all sHsps tested (see Tab. 1), changing from 0.31±0.01 to 0.35±0.01 and 0.34±0.02 for HspB1 and HspB4, respectively. The increase in anisotropy of FUS for the dilute and dense phase indicates slower rotation of FUS molecules in the presence of sHsps, due to molecular interactions between both molecular species. Whereas, when we looked at the anisotropy change of HspB1 after the addition of FUS, we can observe an overall decrease of anisotropy (see Fig. 2 C), shown by the dilute phase anisotropy for HspB1 only at 0.32±0.01 and in the presence of FUS at 0.30±0.01 and 0.29±0.01 for dilute and dense phase, respectively (see Tab. 1). Those lower values likely originate from smaller HspB1 and thus more mobile complexes, most likely by dissociation of the large HspB1 constitutions to smaller molecules, initiated by FUS interactions. Furthermore, we were able to distinguish the anisotropy between the core and the interface of HspB1 in FUS droplets by thresholding, indicating an embedding of HspB1 in FUS droplets by molecular interactions. By looking at the anisotropy resolved image of HspB1 in the presence of FUS (see Fig. 2 B bottom), a dip in anisotropy directly at the droplet interface can be observed, much lower than the anisotropy values observed inside or outside the FUS droplets. This indicates a higher mobility of HspB1 at this location and indicates disassembly of the chaperone complex.

**Fig. 2.**
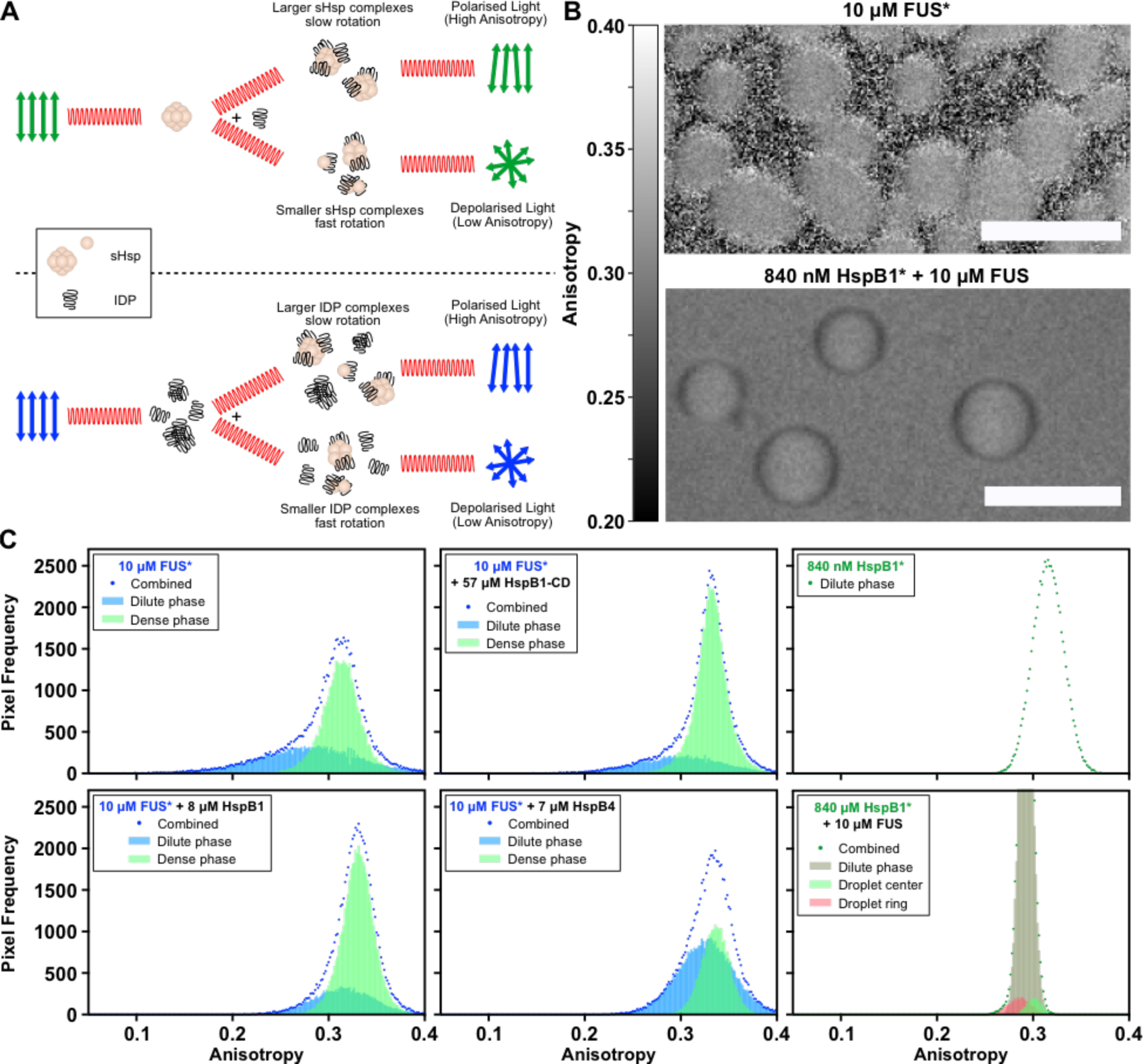
Anisotropy measurements of FUS and sHsps. **(A)** Schematic representation of anisotropy measurements showing excitation of chaperones (top row) or FUS (bottom row) with polarised light. Depending on the formation of larger complexes and thus slow rotation or smaller complexes or even dissociation leading to fast rotation, respectively. **(B)** Pixel-by pixel anisotropy heatmap of confocal scans of 10 µM FUS (top) and of 840 nM HspB1 in the presence of 10 µM FUS (bottom). Scale bar represents 5 µm. **(C)** Anisotropy distributions of various FUS and sHsp combinations. Wherever MBP-FUS-mscarlet3 was used, it was supplemented in a 1:500 ratio. All chaperones were tagged with meGFP. * represents the molecule which was excited.

**Table 1:**
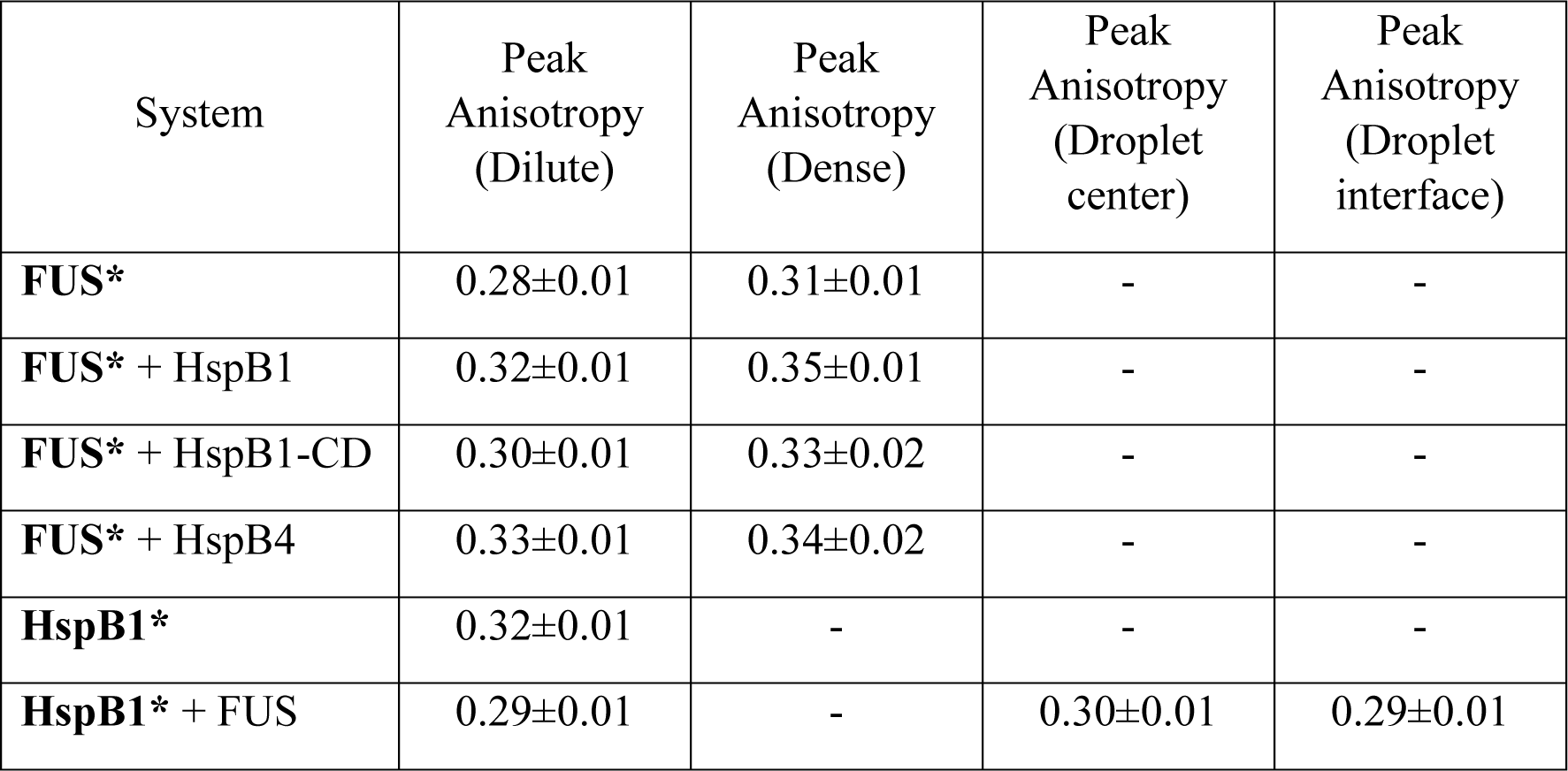
Minimum, peak and maximum anisotropy values of the dense and dilute phase of various FUS-sHsp combinations, respectively. Values were obtained from Fig. 2.

| System | Peak Anisotropy (Dilute) | Peak Anisotropy (Dense) | Peak Anisotropy (Droplet center) | Peak Anisotropy (Droplet interface) |
| --- | --- | --- | --- | --- |
| **FUS\*** | 0.28±0.01 | 0.31±0.01 | - | - |
| **FUS\* + HspB1** | 0.32±0.01 | 0.35±0.01 | - | - |
| **FUS\* + HspB1-CD** | 0.30±0.01 | 0.33±0.02 | - | - |
| **FUS\* + HspB4** | 0.33±0.01 | 0.34±0.02 | - | - |
| **HspB1\*** | 0.32±0.01 | - | - | - |
| **HspB1\* + FUS** | 0.29±0.01 | - | 0.30±0.01 | 0.29±0.01 |

### Effect of small heat shock proteins on disordered proteins below saturation concentrations

Recent developments in the field of microfluidics further enable the study of multiple orthogonal molecules within a heterogeneous solutions, simultaneously and thus, trace specific molecular interactions in a complex environment ^24,47–50^. In order to follow the molecular interactions of sHsps below the saturating concentrations of FUS, we applied microfluidic diffusional sizing, allowing the absolute and simultaneous quantification and sizing of both interaction partners from monomers to nano-clusters in a time-resolve manner. Using this application, we were able to follow the change in hydrodynamic radius of specific sHsps and FUS individually over time. This is achieved by recording lateral diffusion profiles of labelled molecules at different positions along a microfluidic chip (see Fig 3 A). The acquired diffusion profiles were fitted to model simulations on the basis of advection-diffusion equations assuming a monomodal or bimodal Gaussian distribution^51^. In the case of a bimodal distribution, the individual radii, and the quantity of the individual species could be determined from the area under the curves of the two Gaussian populations.

As previous investigations have already shown, FUS can form nano-sized protein cluster (see Fig. 3 B-D)^38,39^. Our protein construct, consisting of an N-terminal maltose-binding protein (MBP) fused to wild-type FUS, should be soluble, even at concentrations where FUS phase separation would be expected. However, by adding TEV protease to the system, the solubility tag is released and molecular interactions and nano-cluster formation is initiated. This way, we were able to probe the interaction of monomeric/dimeric FUS in the presence of HspB1, HspB1-CD or HspB6, respectively, and their influence of FUS oligomerization in a controlled fashion (see Fig. 3 B-D). For all sHsps tested, a concentration dependent inhibition of nano-cluster formation can be observed after the MBP-tag was released. Concentrations below 7.4, 6.6 and 17.4 µM HspB1, HspB1-CD and HspB6, respectively, where sufficient to abolish the formation of detectable nano-clusters completely. However, 1.4, 0.7 and 1.7 µM of the individual chaperones were not enough to interfere with the emergence of FUS cluster at a concentration of 0.5 µM FUS. The inhibition of FUS nano-cluster seems therefore independent of initial sHsp conformation, as the oligomeric HspB1 but also the monomeric/dimeric HspB1-CD and HspB6 can prevent such formation. Furthermore, it seems that sHsps needs to be present in molar excess compared to FUS. Using microfluidic diffusional sizing, we could show that sHsps are capable to control the formation of nano-clusters in a concentration dependent manner, present below saturation concentration. Another observation we can make from the diffusional sizing is a very minor increase, if at all, of molecular size of sHsps in the presence of FUS and FUS nano-clusters. This indicates that the chaperones interact rather with monomeric and dimeric FUS than with FUS nano-clusters. The significance of this preferential binding becomes even more compelling when considered alongside our finding that higher concentrations of sHsps led to smaller droplet sizes (see Suppl. Fig 2), independent on the chaperone size. This change in droplet size may, in fact, be a consequence of the sequestration of FUS monomers by sHsps.

**Fig. 3.**
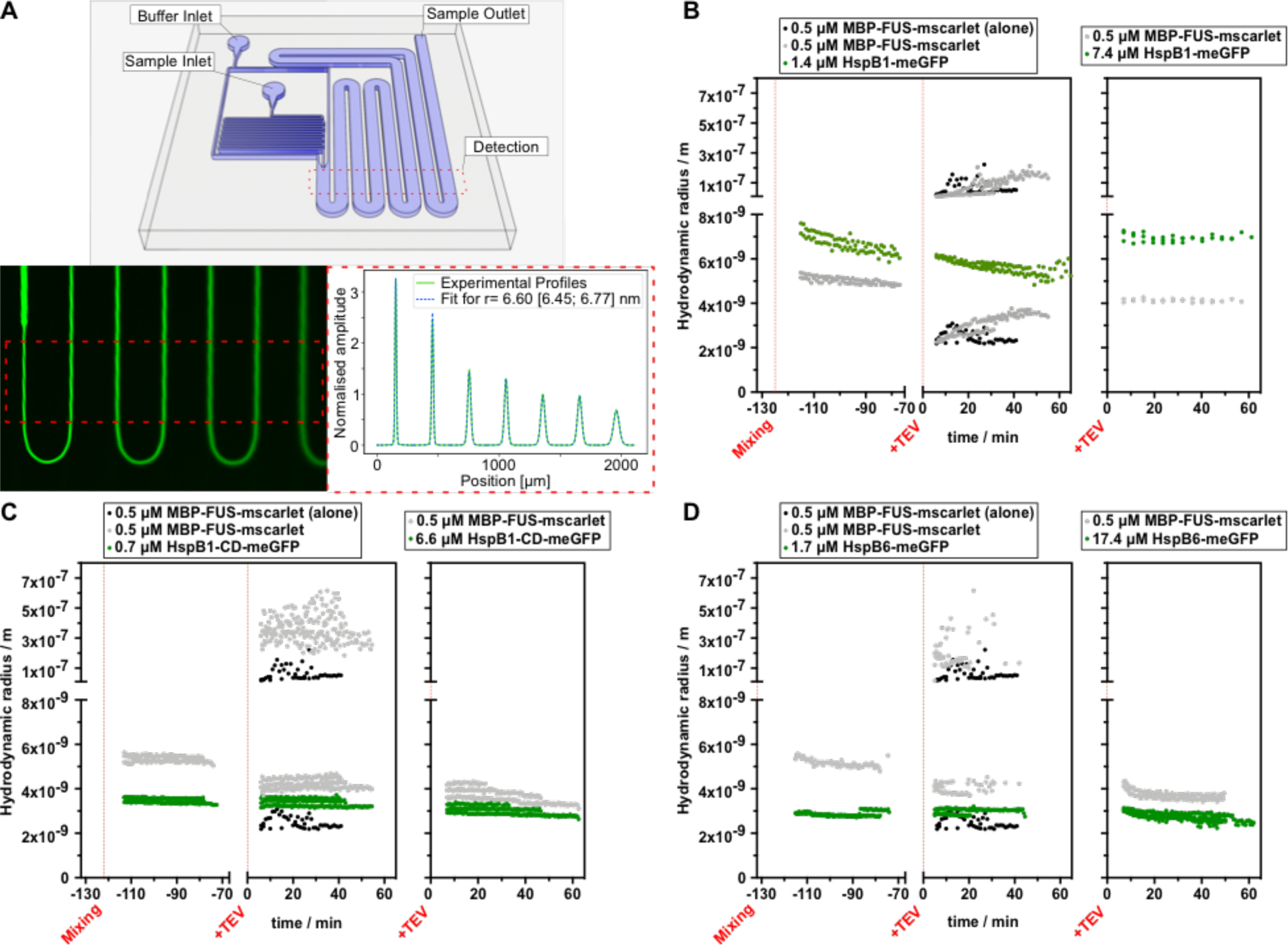
Microfluidic diffusional sizing of sHsps together with FUS. **(A)** Schematic diagram of the microfluidic diffusional sizing device used in this work indicating its most relevant components (top). On the bottom left is an exemplified fluorescence microscope image showing the snakelike channels along the microfluidic device. On the right are the corresponding diffusion profiles (green curve) with the best fit to simulated profiles (dashed blue curve). For this example, the best fit aligns best with the diffusion profile of a molecule with a hydrodynamic radius of 6.6 nm [6.45 nm; 6.77 nm] **(B-D)** Kinetic diffusional sizing before and after the addition of TEV of 0.5 µM MBP-FUS-mscarlet3 only (black dots). Furthermore, each plot shows an overlaid size kinetic of 0.5 µM MBP-FUS-mscarlet3 (grey dots) in the presence of 1.4 µM and 7.4 µM HspB1-meGFP (green dots in B), 0.7 µM and 6.6 µM HspB1-CD-meGFP (green dots in C) and 1.7 µM and 17 µM HspB6-meGFP (green dots in D), respectively. For simplicity, only one pre mixing data set is shown per experimental condition in B, C and D, as those are similar to their respective sHsp experiments without TEV.

## Discussion

Our study delves into the complex interplay between two inherently dynamic systems: IDPs, exemplified by FUS, and sHsps. On one hand, IDPs like FUS exhibit a wide range of behaviours, existing as monomers, undergoing phase separation to form larger clusters, and even aggregating into aberrant forms^13,15,39,52^. On the other hand, sHsps present their own dynamic nature, forming complexes of varying sizes and localizing preferentially at certain cellular sites^43,53,54^. Interestingly, a similar disassembly process of sHsps upon interacting with FUS, as we can see for HspB1 in our anisotropy data, has been reported for HspB5 binding to α-synuclein fibrils^24^. In this case, the disassembly of the sHsp has been shown to provide the binding energy for chaperone binding to fibrils by entropic contribution. The entropic gain was hypothesised to originate from the chaperone itself through disassembly of the sHsp complexes, and not the solvent.

These protein clusters, tens to hundreds of nanometres in size, are multimers, but differ from aggregates or protein fibrils, due to their still dynamic not thermodynamically arrested state and have been reported to be present below and above protein saturation concentration^38,39^. Within this context, we observed that the presence of sHsp exerted a strong inhibitory effect on the formation of FUS nano-clusters. This observation adds a layer of complexity to our understanding of how sHsps might intervene in the varied behaviours of IDPs, potentially offering a mechanism to regulate or even suppress undesirable phase transitions and aggregations. Nevertheless, whether such nano-clusters share similarities to off- or toxic on-pathway oligomers known from amyloid formation is unclear^7,55^, as well as whether they serve as catalytic seeds for cluster formation and phase separation of similar or orthogonal biomolecules.

Further enriching this narrative is our finding that larger sHsp complexes preferentially accumulate at the interface of prefibrillar droplets. This location-specific behaviour contrasts sharply with that of monomeric sHsp units, which were notably absent at these interfaces. This is of particular interest, as recent studies indicate that the phase between the dense and dilute phase of droplets is the source for fibril formation or protein aggregation^12,13,15,42^.

The dynamic interplay between IDPs and sHsps (see Fig. 4), each with their own complex behaviours, opens new avenues for understanding how cells might finely tune their response mechanisms to prevent pathological outcomes often linked to neurodegenerative diseases.

**Fig. 4.**
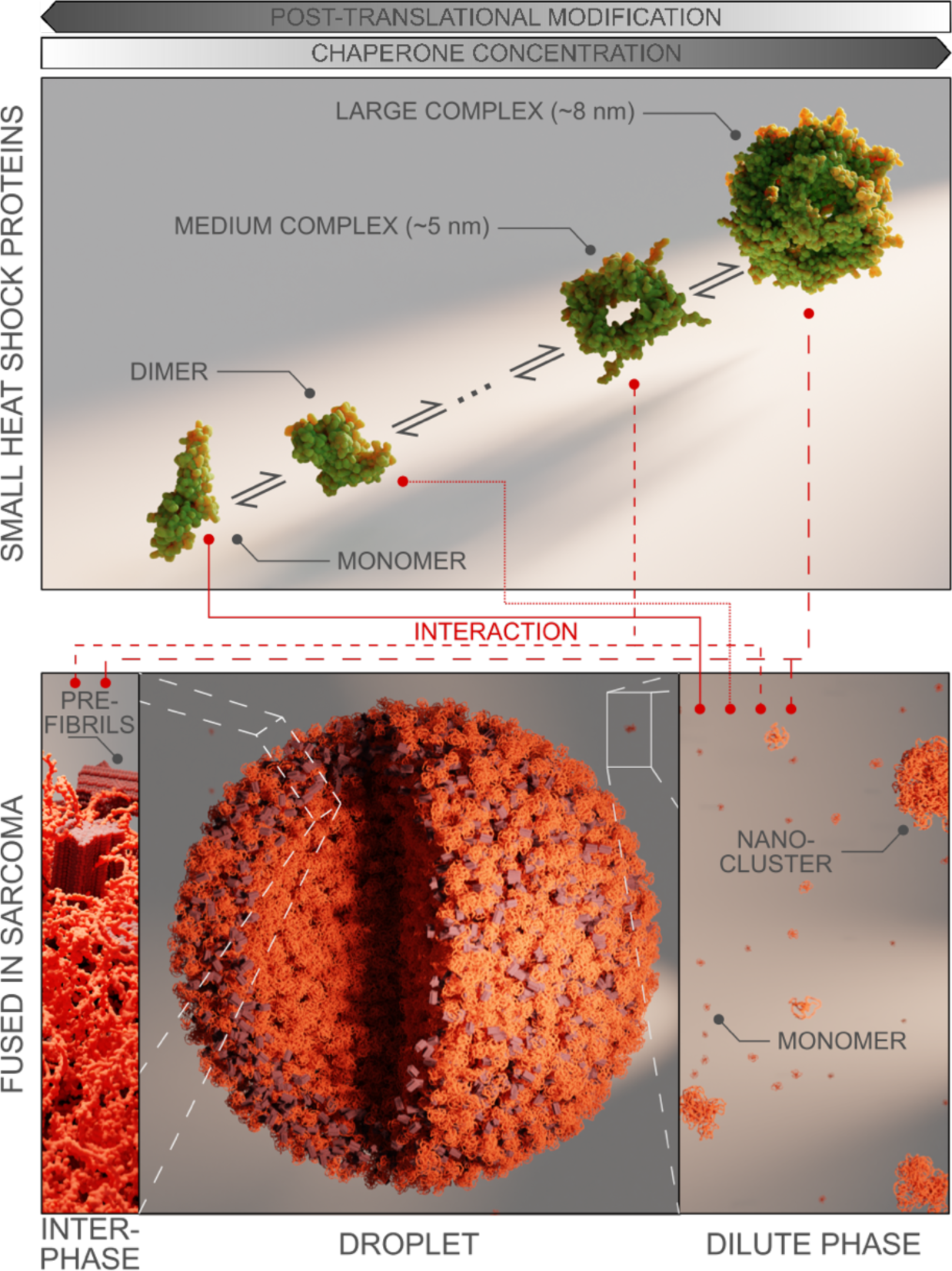
Schematic summary of dilute phase and dense phase composition found for many proteins and how sHsps interplay with such components. Proteins can phase separate into densely packed droplets, where for FUS it has been shown that protein aggregation is promoted at the dense and dilute protein interface. The dilute phase on the other hand, is composed of freely diffusing monomers, oligomers or even nano-clusters (bottom row). The diverse components are accessed individually by specific components of the dynamic sHsp system. The abundance of those specific sHsp components is tightly regulated by post-translational modifications or overall protein concentration (top row). Images were rendered using the HspB1 (PDB: 6DV5), FUS alpha fold (PDB: AF-P35637-F1) and FUS fibril (PDB: 7VQQ) structures.

Future work could extend these findings by exploring how sHsps interact with more complex droplets composed of multiple IDPs and other biomolecules. Techniques like microfluidics could offer additional perspectives, particularly in understanding the thermodynamics underlying these interactions.

In sum, our research contributes to a growing body of knowledge about the prevention of dynamic arrest leading to pathological liquid-to-solid transitions with the potential to allow the development of future therapeutic strategies aimed at mitigating the risks associated with pathological protein aggregation or even restoration and regulation of condensates or nano-clusters in the first place in the context of neurodegenerative diseases.

## Methods

### Protein expression and purification

MBP–FUS–His_6_ wt, MBP–FUS–mscarlet3-His_6_ wt^56^ and TDP-43-MBP-His_6_ wt^57^ were expressed and purified by adapting previous protocols. In short, FUS was transformed into BL21-DE3-Rosetta-LysS and bacteria were grown in LB medium at 37°C to an absorbance at 600 nm of 0.8. Expression was induced with 1 mM IPTG for 22 h at 12°C. Cells were lysed in lysis buffer (50 mM sodium phosphate, pH 8.0, 300 mM NaCl, 40 mM Imidazole, 10 µM ZnCl_2_, 4 mM β-mercaptoethanol (BME), 10% glycerol) using sonication. After centrifugation, the supernatant was further purified using Ni-NTA beads and washed with lysis buffer. The protein was eluted with elution buffer (50 mM sodium phosphate, pH 8.0, 300 mM NaCl, 250 mM Imidazole, 10 µM ZnCl_2_, 4 mM BME) and added to an amylose resin (New England Biolabs), incubating overnight at 4°C. The mixture was further supplemented with an equal amount of salt-free buffer (50 mM sodium phosphate, pH 8.0, 40 mM Imidazole, 10 µM ZnCl_2_, 4 mM BME) and kept mixing for another 1 h. Finally, the amylose resin mixture was transferred to a column and washed with amylose washing buffer (50 mM sodium phosphate, pH 8.0, 300 mM NaCl, 40 mM Imidazole, 10 µM ZnCl_2_, 4 mM BME) and eluted with amylose elution buffer (50 mM sodium phosphate, pH 8.0, 300 mM NaCl, 40 mM Imidazole, 10 µM ZnCl_2_, 20 mM maltose, 4 mM BME).

TDP-43 was expressed in BL21-DE3-Rosetta 2 cells in LB at 37°C by adding 1 mM IPTG when OD has reached 0.6 and kept shaking overnight at 12°C. Cells were lysed in lysis buffer (20 mM Tris, pH 8.0, 1 M NaCl, 10 mM imidazole, 4 mM BME, 10% glycerol, 1 µg/ml each of aprotinin, leupeptin and pepstatin) together with 100 µg/ml lysozyme and RNase A and further sonication. After centrifugation the supernatant was incubated with Ni-NTA beads and washed with lysis buffer and eluted with elution buffer (20 mM Tris, pH 8.0, 1 M NaCl, 300 mM imidazole, 4 mM BME, 10% glycerol, 1 µg/ml each of aprotinin, leupeptin and pepstatin). Eluted product was loaded on a HiLoad 16/60 Superdex 200 prep grade column (GE Healthcare) and run with a sample buffer (50 mM Tris, pH 8.0, 300 mM NaCl, 5% glycerol, 1 µg/ml, 2 mM TCEP).

The small heat shock proteins HspB1-meGFP-His_6_, HspB4-meGFP-His_6_, HspB5-meGFP-His_6_, HspB6-meGFP-His_6_, as well as the N- and C-terminus deficient HspB1-core domain (HspB1-CD-meGFP-His_6_) were expressed with BL21-AI cells in LB. When OD reached 0.6, the expression was induced by the addition of 1 mM IPTG and 0.02% (w/v) arabinose and kept shaking overnight at 18°C. Cells were lysed in lysis buffer (20 mM Tris, pH 7.9, 100 mM NaCl, 5 mM imidazole, 0.1 mM PMSF) using a continuous flow cell disruptor (Constant Systems Ltd.). The lysate was centrifuged at 20.000xg for 60 min and the supernatant was further incubated with Ni-NTA beads for 2 h at 4°C. The Ni-NTA beads were loaded on a column and washed using the lysis buffer with 20 and 30 mM imidazol, respectively. The protein was eluted with lysis buffer containing 100, 200, 300 and 400 mM imidazol. Relevant fractions were combined and run over size-exclusion (HiLoad S200 16/60 prep grade, GE Healthcare) with sample buffer (20 mM Tris, pH 7.9, 100 mM NaCl, 1 mM DTT, 10% Glycerol)

TEV protease was expressed in BL21-RIL cells. Therefore, cells were grown at 37°C and 1 mM IPTG was added add mid log-phase. After induction, the temperature was reduced to 30°C and left shaking for 4 h. Cells were lysed in lysis buffer (50 mM sodium phosphate, pH 8.0, 100 mM NaCl, 25 mM Imidazole, 10% glycerol) using a continuous flow cell disruptor (Constant Systems Ltd.). Lysate was centrifuged at 15.000xg for 30 min. Supernatant was filtered through a 0.25 µm membrane and loaded onto a pre-packed Ni-NTA column. Sample was washed with lysis buffer and eluted with an increasing gradient of elution buffer (50 mM sodium phosphate, pH 8.0, 100 mM NaCl, 200 mM Imidazole, 10% glycerol) supply. Eluted sample was buffer exchanged to 25 mM sodium phosphate, pH 8.0, 200 mM NaCl, 2 mM EDTA, 10% glycerol and concentrated to 2 mg/ml.

All chemicals were of analytical grade and purchased from Sigma Aldrich unless otherwise stated.

### Droplet assays for microscopy

Purified full length MBP-FUS or TDP-43-MBP were diluted in droplet buffer, which was 10 mM HEPES pH 7.2, 200 mM KCl, 3% glycerol, 1 mM DTT and 20 mM HEPES pH 7.5, 150 mM NaCl, 1 mM DTT buffer, respectively. MBP-FUS was supplemented 1:100 with MBP-FUS-mscarlet3. Furthermore, individual sHsps were added if stated. Phase separation was induced by the addition of TEV to a final concentration of 0.1 mg/ml. Images were taken with a BC43 spinning disk confocal (Andor Technology Ltd.).

For the linescan intensities the background intensity was determined from buffer blanks and subtracted for both channels and the intensity was further normalised to the maximum intensity of the individual scans.

In order to follow the reduction of droplet size in dependence of chaperone concentration, nine tiles of 121 µm x 125 µm, each, after about 30 min of incubation. Those tiles were analysed using a python script. The code automates particle size analysis in microscopy images by applying a threshold to identify particles, measures their sizes, filters based on predefined criteria like circularity (between 0.7 and 1) and minimum area, and provides an average size.

### FRAP of droplets and analysis

Droplets were prepared as described above. The meGFP signal of HspB1 for individual droplets was bleached. The used setup was a Stellaris 8 Falcon confocal microscope (Leica Microsystems GmbH). The 488-laser intensity for bleaching was 50% with 0.5 ms/px reaching about 500-700 ms pulse. Droplet movement was drift corrected using the Fast4DReg plugin in Fiji^58^. Fluorescence Recoveries of droplets were baseline corrected and normalised to lowest and highest intensity after and before bleaching, respectively. We employed a Python script for image analysis on the FRAP image stacks. The process involved loading images, applying thresholding to identify regions, and calculating region-specific statistics (average intensity, standard deviation) across all frames. Later on, the fluorescence recovery was fitted to a two-phase association model.

### Anisotropy measurements and analysis

For anisotropy measurements droplet assays were performed as described. However, for the anisotropy measurements 12.5 nM MBP-FUS-meGFP was added to 10 µM MBP-FUS solution. Depending on the individual experiment sHsps were present. Phase separation was induced by the addition of TEV. For the measurements a self-build confocal microscope setup, described before^59^, with a 50 µm pinhole was used. In short, samples were excited with a picosecond pulsed laser diode head including the wavelengths of 485 nm (LDH-D-C-485, PicoQuant) (8% power) to measure HspB1 anisotropy, 560 nm (LDH-D-TA-560, PicoQuant) (35% power) to measure FUS anisotropy. The beam travelled through a Glan-laser polarizer (Thorlabs) set to vertical position and was directed into a laser scanning system (FLIMbee, PicoQuant). The three galvo mirrors in the scanning system were imaged onto the backfocal plane of the objective (60x SR Plan Apo IR, 1.27 NA, Nikon). The dwell time was 150 µs/px and a frame size of 256x256 px. For each final image, at least 380 frames were acquired, and the intensity was added up. This results in two images, one for parallel and rectangular emission.

A python script was used for image evaluation. The code analyses the global images or applies an intensity threshold before to separate dense and dilute phase of the droplets. The applied analysis, however, is the same. For each pixel we apply the following equation:

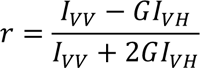

where r is the fluorescence anisotropy, *I_vv_* corresponds to the vertical polarised excitation and vertical polarised emission, *I_vH_* corresponds to the vertical polarised excitation and horizontal polarised emission and the G factor^60^. The images were on one hand converted into an anisotropy heat map with each pixel giving its individual anisotropy value and on the other hand, distributions of the anisotropy measurements were plotted for all analysed pixels. The distribution for the dense and the dilute area was fitted to a normal distribution and the mean with error is given in Table 1. In order to calculate the average photon count per pixel for the same images, the equation:

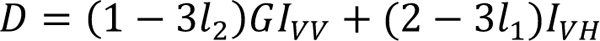

is applied to each individual pixel in a region of interest, with D being the photon count, *l*_1_ and *l*_2_ are factors accounting for polarization mixing caused by the high numerical aperture objective lens. All values are average for the entire area of interest.

### Fabrication and use of microfluidic diffusion devices

The fabrication and the operation of the microfluidic diffusion device used in the present studies have been described in previous papers^51,61^. Briefly, the microfluidic chips were fabricated in PDMS by using standard soft lithography. The sample to be analysed and the buffer were introduced into the system through reservoirs connected to the inlets, and the flow rate in the channel was controlled by applying a negative pressure at the outlet by a syringe pump (Cetoni neMESYS, Korbussen, DE); at typical flow rates in the range from 160 µl/h to 200 µl/h. Two different devices were used. For one, lateral diffusion profiles were recorded at four different positions (1.2 mm, 10.2 mm, 20.2 mm and 40.2 mm) for the other, 7 different positions (1.0 mm, 1.6 mm, 10.3 mm, 10.9 mm, 19.5 mm, 20.1 mm and 38.2 mm) by using a Visitron spinning-disk microscope (Visitron Systems GmbH) together with an ORCA-flash 4.0 camera C13440 CMOS camera (Hamamatsu Photonics). Samples were added to the microfluidic chip via a tip reservoir for continuous measurements, either already mixed with an individual chaperone or without. At the same time, an aliquot was kept at room temperature to have consistent conditions. After about two hour, TEV protease was added to this aliquot to a final concentration of 0.1 mg/ml and added to a new chip, again for continuous measurements. The TEV protease cleaves off the MBP-tag and thus initiates cluster formation.

The diffusion profiles were fitted to numerical model simulations based on advection-diffusion equations for mass transport under flow^51^. Either an average monomodal or a bimodal distribution were assumed. For the latter, from the area under the curves of the two Gaussian populations, the concentrations of each individual component were evaluated.

## Acknowledgments

We thank all of the members of the Lemke and Dormann laboratory for helpful discussions; the core facilities of the Faculty of Biology at Johannes Gutenberg University Mainz; and the Protein Production Core Facility at the Institute of Molecular Biology Mainz for expert assistance. T.S. was funded by the EMBO Postdoctoral Fellowship (ALTF 1020-2020). E.A.L. acknowledges funding from the CRC1551 ‘Polymer concepts in cellular function’ of the Deutsche Forschungsgemeinschaft (DFG project number 464588647) and SPP2191 (DFG project number 419070619).

## Author contributions

T.S. and E.A.L. designed research; T.S. performed research; T.S. and E.A.L. analysed data; and T.S. and E.A.L. wrote the paper.

