## Supplementary information for "Chaperone quality control in liquid-phase separated organelles"

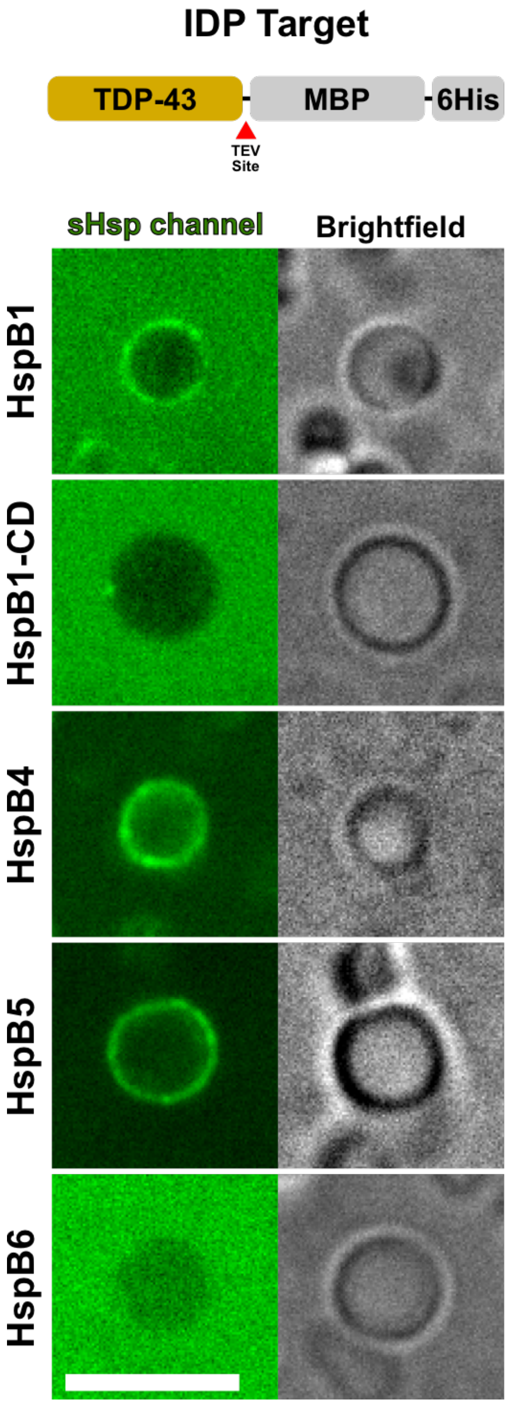


**Suppl. Fig. 1. Effect of small heat shock proteins on TDP-43 above saturation conditions.** (Top) Schematic of used protein constructs with TDP-43 as another representative IDP target. Wild-type TDP-43 construct contains a C-terminal maltose-binding protein (MBP) for increased solubility with an additional C-terminal 6-histidine tag. Both can be cleaved off by TEV protease. Below, spinning disk images of bulk droplet assays with 10 µM TDP-43 and 3 µM HspB1, 5 µM HspB1-CD, 5 µM HspB4, 6 µM HspB5 and 9 µM HspB6 in 20 mM HEPES pH 7.5, 150 mM NaCl, 1 mM DTT buffer, respectively (left). Phase separation was triggered by the addition of TEV. Scale bar is 5 µm.

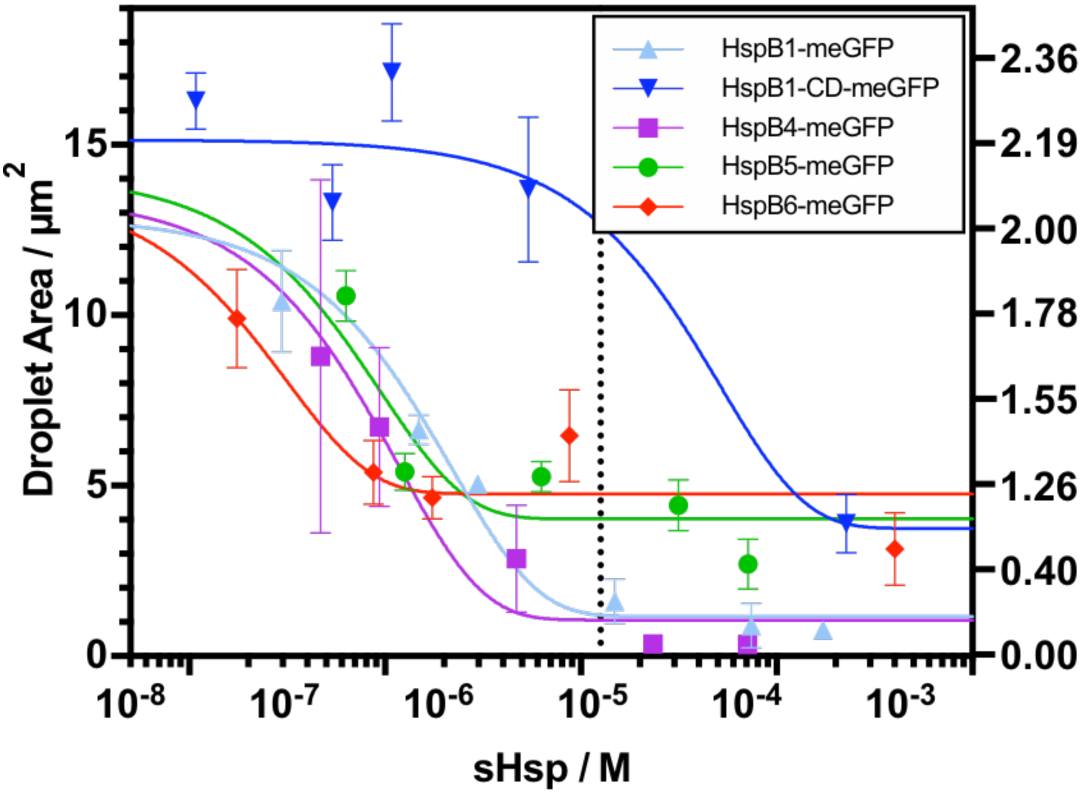


**Suppl. Fig. 2. FUS droplet area and corresponding radius in dependency of sHsp concentration present.** Spinning disk images of bulk droplet assays with 10 µM FUS and various sHsp concentrations were performed in 10 mM HEPES pH 7.2, 200 mM KCl, 3% glycerol, 1 mM DTT buffer. FUS sample was supplemented with 1:100 MBP-FUS-mscarlet3 and phase separation was triggered by the addition of TEV. After about 30 min 9 tiles of 121 µm x 125 µm, respectively, were taken and used for further evaluation. For determining the average droplet area, an intensity threshold was applied with further circularity and area limits on all images.

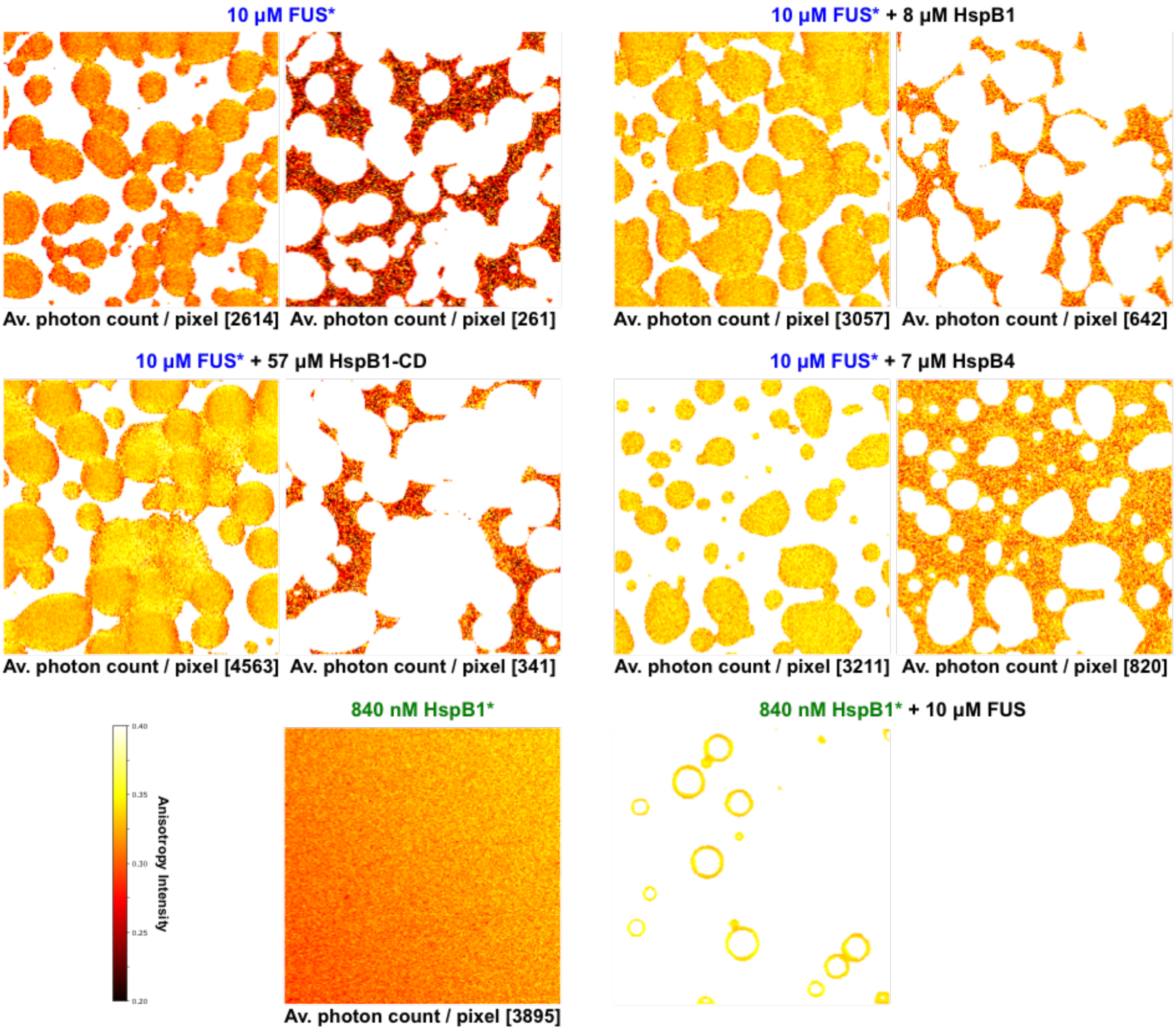


**Suppl. Fig. 3. Pixel-by-pixel anisotropy heatmap of confocal scans of dense and dilute phase ROIs corresponding to Fig 2.** In the presence of FUS, phase separation was induced by the addition of TEV and the release of the MBP-tag. * represents the molecule, which was excited, which was in case of FUS an mscarlet3-tag and in case of HspB1 an meGFP-tag. Each frame has a dimension of 26 µm x 26 µm.

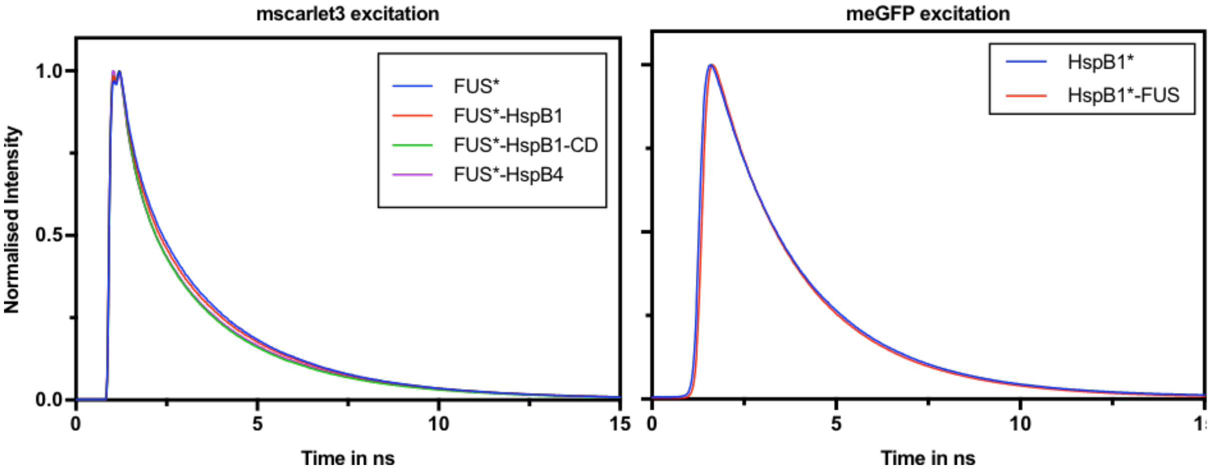


**Suppl. Fig. 4. Lifetime curves of the measurements of Supplementary Figure 3.** For each condition, intensity decays of accumulated frames of both the polarised emissions, vertical and horizontal, were summed up and normalised to overall highest intensity. For FUS, the mscarlet3-tag was excited and for HspB1, the meGFP-tag was excited.

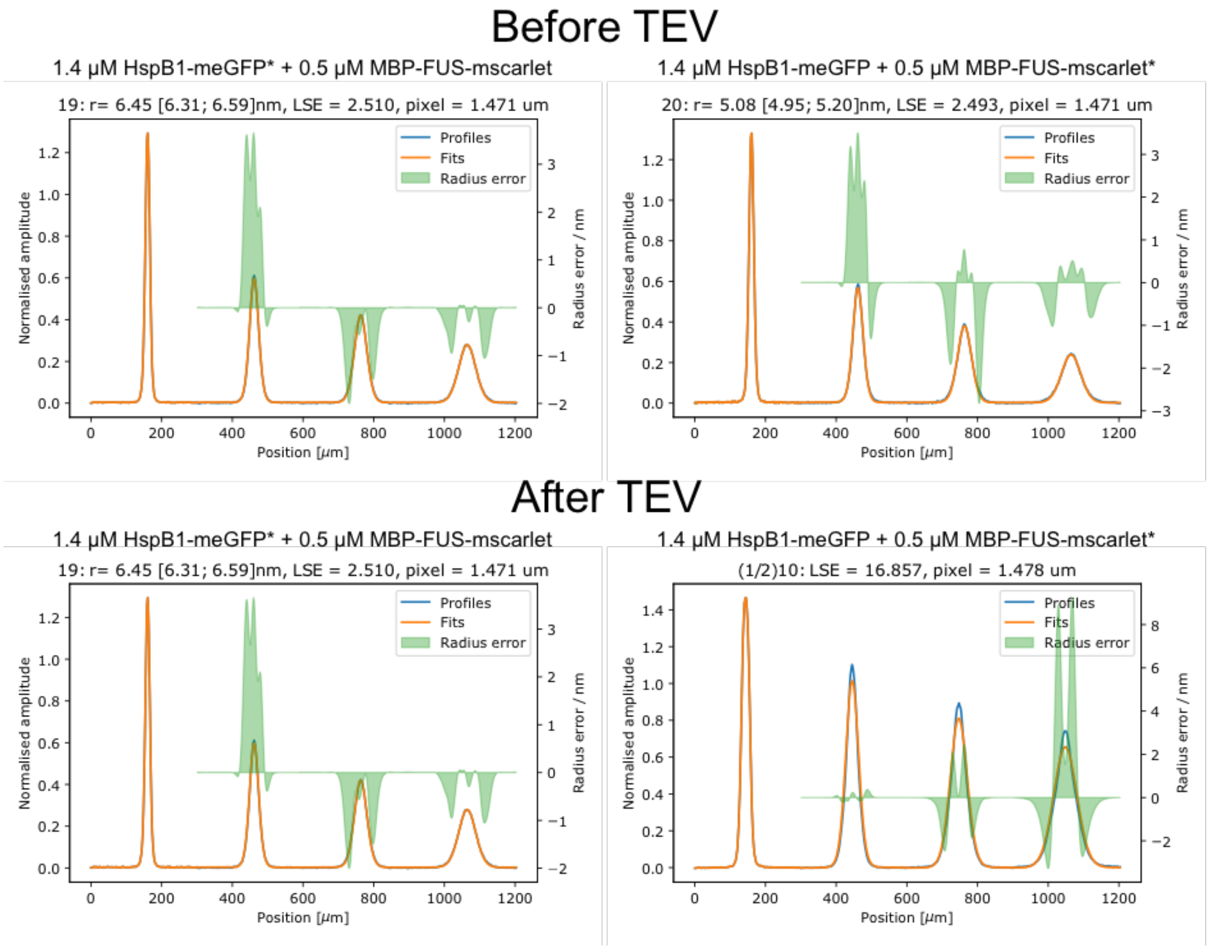


**Suppl. Fig. 5. Example diffusion profiles of kinetic measurements seen in Figure 3B.** Top row shows the trace fittings of 1.4 µM HspB1-meGFP (left) and 0.5 µM MBP-FUS-mscarlet3 (right), run and measured simultaneously, before adding TEV. The bottom row shows the same after adding TEV.
